# Electrochemical Detection of Adrenaline and Hydrogen Peroxide on Carbon Nanotube Electrodes

**DOI:** 10.1101/2021.08.24.457486

**Authors:** Gaurang Khot, Mohsen Kaboli, Tansu Celikel, Neil Shirtcliffe

**Affiliations:** Department of Neurophysiology, Donders Institute for Brain, Cognition and Behavior, Radboud University, The Netherlands; Faculty of Technology and Bionics, Hochschule Rhein-Waal University of Applied Sciences, Kleve, Germany; Robotics, Tactile and Artificial Intelligence Bayerische Motoren Werke AG (BMW), Munich, Germany

**Keywords:** FSCV, HYDROGEN PEROXIDE, ADRENALINE, CNT, CYCLIC VOLTAMMETRY, ELECTROCHEMISTRY, NANOTUBES

## Abstract

Adrenaline and hydrogen peroxide have neuromodulatory functions in the brain. Considerable interest exists in developing electrochemical sensors that can detect their levels *in vivo* due to their important biochemical roles. Challenges associated with electrochemical detection of hydrogen peroxide and adrenaline are that the oxidation of these molecules usually requires highly oxidising potentials (beyond 1.4 V vs Ag/AgCl) where electrode damage and biofouling are likely and the signals of adrenaline, hydrogen peroide and adenosine overlap. To address these issues we fabricated pyrolysed carbon electrodes coated with oxidised carbon nanotubes (CNTs). Using these electrodes for fast-scan cyclic voltammetric (FSCV) measurements showed that the electrode offers reduced overpotentials compared with graphite and improved resistance to biofouling. The Adrenaline peak is reached at 0.75(±0.1) V and reduced back at -0.2(±0.1) V while hydrogen peroxide is detected at 0.85(±0.1) V on this electrode. The electrodes are highly sensitive with a sensitivity of 16nA *µ*M^-1^ for Adrenaline and 11nA *µ*M^-1^ for hydrogen peroxide on an 80 µm^2^ electrode. They are also suitable to distinguish between adrenaline, hydrogen peroxide and adenosine thus these probes can be used for multimodal detection of analytes.

## Introduction

Fast Scan Cyclic Voltammetry (FSCV), where the voltage sweep is extremely fast, has received considerable attention for the detection of neurotransmitters and neuromodulators as it increases sensitivity and measurement speed (1,2). The ability of FSCV to detect nanomolar concentrations of a target analyte using chemically modified electrodes (3) is of particular interest to neuroscientists as it allows the detection of physiologically relevant concentrations of various bioactive substances in the living brain (2). Challenges associated with the development of chemically modified electrodes as brain implants are the degradation of chemical probes over time, loss of sensitivity, overlapping signal from analytes, biofouling and immune responses at the site of implantation (2). These issues have limited the application of carbon fibre electrodes, which are considered to be the gold standard material for *in vivo* and *in vitro* preclinical applications because other conventional materials have even less favorable properties (2). Moreover, carbon fibre electrodes commonly cannot resolve dopamine and serotonin (4), hydrogen peroxide and adenosine from one another (2,5,6), which limits their application in biochemically complex environments like the cerebrospinal fluid.

Considerable interest exists to use carbon nanotubes (CNTs) as an alternative electron carrier. Thanks to their small size, faster electron transfer kinetics, reduced overpotentials, biostability and resistance to biofouling CNTs have become an attractive material for the electrochemical detection of neurotransmitters and neuromodulators (2). Due to their hydrophobic nature and π-bonding, they form aggregates when dispersed in most solvents (7). To address these issues, the functionalization of CNTs can be performed with oxidising acids, which opens the ends of the CNTs and grafts oxygen atoms onto the exposed edge planes making them polar enough to disperse in alcohols and even water(7). This functionalization also allows CNTs to act as a local proton acceptor, thereby reducing overpotential for redox reactions in which proton transfer is required (7). CNT based surfaces have been used in electrochemistry for electrochemical detection of neurotransmitters and neuromodulators (8–12).

Neuromodulators are an important class of compounds that manipulate neuronal activity, e.g. by altering the firing pattern of neurons (1). Adrenaline is a monoamine neurotransmitter that serves a dual purpose, both as a hormone and a neurotransmitter (13). In the brain, adrenaline is responsible for various cognitive functions, including alertness and flight or fight response (14) Dysregulation of adrenaline is known to cause depression and anxiety (14).

Hydrogen peroxide, superoxide(•O_2_) and hydroxyl radicals(•OH) are generated as the end product of energy metabolism and are usually considered to be waste/toxic materials to cells. Recent evidence suggests that peroxide molecules also play a significant role in cellular signalling and information passing, similar to neurotransmission (15–17). The dual nature of hydrogen peroxide as a neuromodulator and in energy generation makes it attractive to be able to measure.

Electrochemical detection of adrenaline on carbon electrodes is reported at a highly anodic potential above +1.3 V (13,18). The use of such highly oxidising potential causes degradation of electrodes and generation of reactive oxygen species (ROS) at the surface of the electrode thereby reducing the sensitivity of electrodes (13). These challenges have limited the ability to measure adrenaline in the brain. Another issue with adrenaline detection in the brain is interference from analytes like hydrogen peroxide and adenosine, which have overlapping signals (1,19,20). Thus the occurrence of identical signals for multiple analytes requires electrode materials on which these analytes can be separated (21). Studies have shown that CNTs have faster electron transfer kinetics than graphite (12), are resistant to biofouling (22) and are able to discriminate between these analytes by shifting the voltages where they occur (4). We therefore exploited these properties of CNTs, by coating them onto pyrolytic carbon electrodes to investigate the electrochemical dynamics of adrenaline and hydrogen peroxide.

## Materials and Methods

Hydrogen peroxide, adrenaline, 4,-(2-hydroxyethyl-)1-piperazineethanesulfonic acid (HEPES), sodium chloride (NaCl), potassium chloride (KCl), sodium bicarbonate (NaHCO_3_), magnesium chloride (MgCl_2_), monosodium phosphate (NaH_2_PO_4_) and Nafion® 1100W (5% in alcohols) were sourced from Sigma Aldrich, Germany. Quartz capillaries (o.d. 1 mm, i.d. 0.5mm, length 7.5 cm) were purchased from Sutter Instruments. Water used in the experiments was deionised in house using a Millipore column and had a resistivity of 18MΩ.

### 1. Fabrication of Microelectrodes

Quartz capillaries (O.D: 1.0mm and I.D: 0.5mm) were pulled using a Sutter puller p-2000 (Sutter instruments). The pulling parameters were: heating temperature 750°C, velocity 50 and DEL 127, which give a tip diameter of 0.8-1.5µm. The tip was broken shorter to reach a final diameter of 25-30µm.

### 2. Pyrolysis

Pyrolysed carbon microelectrodes were fabricated by performing pyrolysis of propane in a nitrogen environment. Quartz capillaries were heated for 1 minute at the narrower end for carbonization. Once the desired duration was reached, the heat was moved towards the broader end. Upon completion of pyrolysis, electrodes were allowed to cool under nitrogen flow.

### 3. Dispersion of CNTs into Nafion Solution

CNTs were functionalized by dispersing them in concentrated hydrochloric acid (HCl) and concentrated sulfuric acid (H_2_SO_4_) in a ratio of (3:1 v:v). The solution was stirred with a magnetic stirrer for 48 hours. Washing was carried out by decanting off the acid and repeated washing and decanting with distilled water until pH 7.0 was reached. After washing, the CNTs were allowed to air dry for 24-36 hours. CNTs were suspended in a solution of Nafion diluted with ethanol and sonicated for 30 minutes prior to use. The suspension was stable for 6—8 weeks. In order to obtain a thin coating of CNTs onto the electrode, the pyrolytic carbon electrodes were dip-coated in this solution for 5 seconds and allowed to dry. The process allowed the deposition of a sufficient amount of CNTs to modify the surface properties of the electrode. Successful coating could be verified by a change in electrochemical signal in a phosphate buffer solution.

### 4. Electrochemical setup

Electrochemical detection of adrenaline and hydrogen peroxide was carried in an electrophysiological setup that had ports for delivery and outflow of solution (23–27). To mimic the rapid change of concentrations observable in the brain a flow cell was set up that delivered the target analyte as a 5-second bolus, followed by a pause of 10 seconds. A constant flow of buffer was maintained at 2mL min^-1^ serving as a background. Electrochemical detection of analytes was carried out using the “Dopamine Waveform” (21), wherein the electrode was cycled from -0.4V to 1.3V and cycled back to -0.4V at 400 V s^-1^ in the presence of a silver wire, coated in AgCl in KCl solution (3.5M) serving as a reference electrode (Ag/AgCl). Data were collected and stored offline for analysis.

## Results and Discussion

### 1. Electrochemical Detection of Adrenaline on Pyrolytic-CNT electrode

The electrochemical behaviour of adrenaline on CNT coated electrodes was studied by introducing 1 *µ*M of adrenaline to an electrode being voltage scanned. The electrode was exposed to 10 consecutive cycles of adrenaline (10 bolus additions) while cycling the potential, initially at 400 V s^-1^. Multiple oxidation peaks and one reduction peak were observed in the cyclic voltammograms at this scan rate. One oxidation peak (0.9V) was associated with oxidation of adrenaline and the second peak was seen at 1.1 V, which was probably associated with biofouling of the electrode and hence we chose to increase the scan rate to 600 V s^-1^ in order to limit this side reaction. Background subtracted voltammograms show that adrenaline is oxidized at 0.75(±0.1) V and the product reduced back at -0.2(±0.1) V (**Figure 1A**). A broad oxidation peak is seen that has a slow rise and fall time, thus suggesting that oxidation of adenosine on pyrolytic-CNT electrodes is a multistep oxidation process occurring on the surface of the electrode.

**Figure 1:**
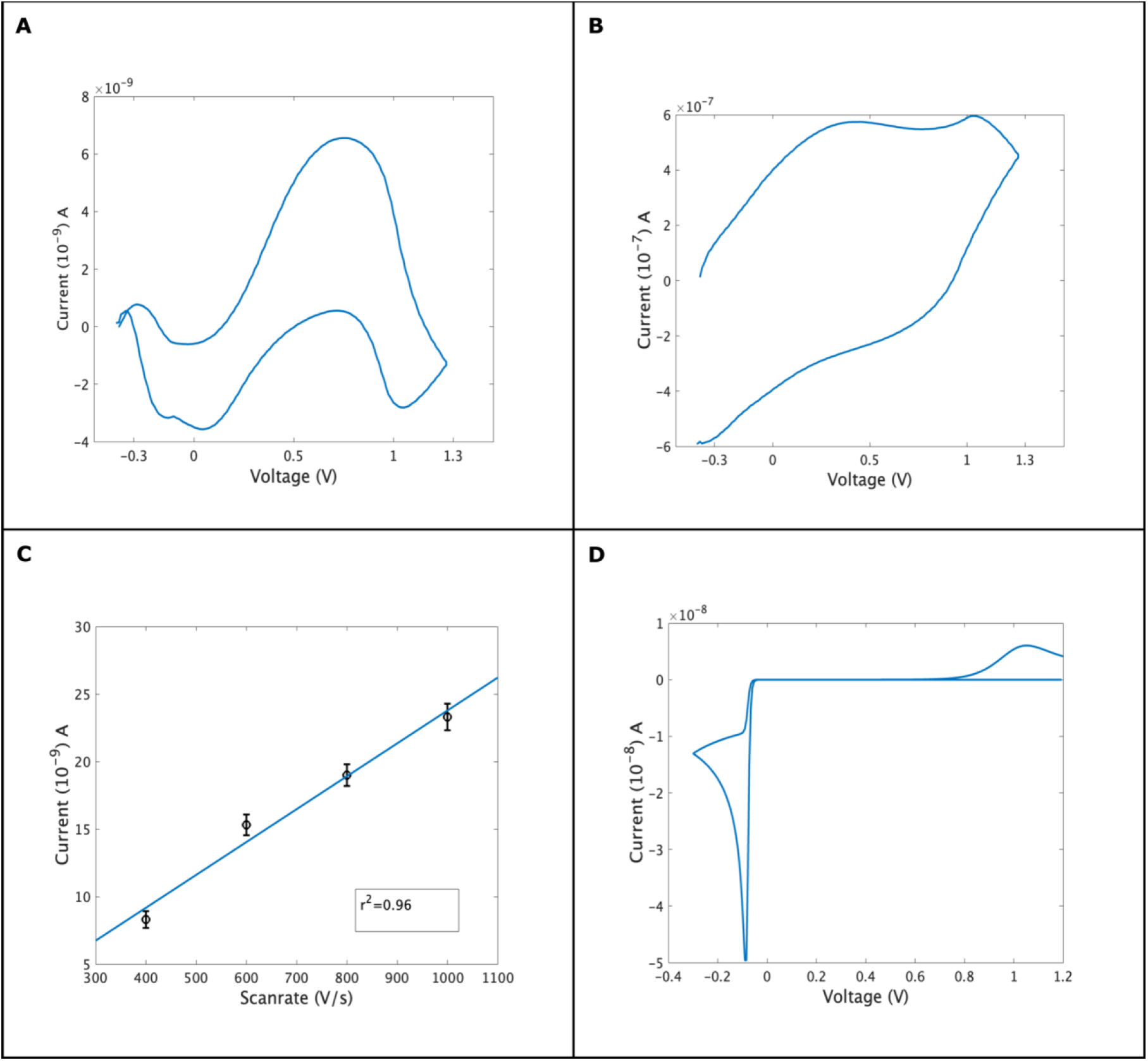
Shows the oxidation of 1 *µ*M adrenaline from -0.4V to 1.3V and cycled back at - adrenaline is oxidized at 0.75(±0.1) V and reduced at -0.2(±0.1) V. **B**: Shows the background current. It should be noted that lysis of water occurs at 1.1 V on CNT electrodes. A reduction peak can be seen at 0.9V. **C**: Shows the peak current is directly proportional to scan rate with a r^2^=0.96 fit, showing the current is dominated by adsorbed species; the electrode has a sensitivity of 16nA *µ*M^-1^ on about 80 *µ*m^2^ **D**: The theoretical prediction for adrenaline, using a diffusion constant of 6×10^−6^ m^2^ s^-1^, the rate of electron transfer was kept at 1 × 10^−6^ m^2^ s^-1^ other parameters are standard.

We further investigated the behaviour of adrenaline on the electrode surface, by varying scan speed and keeping the concentration of adrenaline constant. As the scan rate was varied, peak current increased linearly, suggesting that adrenaline binds to the surface of the electrodes (**Figure 1C**). This type of preadsorption increases the sensitivity of detection by increasing the amount of analyte available to the electrode for a short time, thereby allowing the detection of the nanomolar concentration of the analyte of interest (21, 28). To investigate if this waveform could be used for longer term measurement the electrode was exposed to 10 consecutive cycles of adrenaline for 10 minutes. We did not observe any changes in the voltammogram over this time, suggesting that the CNT surface combined with the increased scan rate limits the polymerization of adrenaline onto CNT electrodes, thus providing long term stability. To understand the oxidation of adrenaline on CNT electrodes, we modelled the rate of electron transfer, diffusion coefficient and chemical rate constant. Mathematical modelling suggests that adrenaline is oxidized at 1.1 V and reduced at - 0.15 V (**Figure 1D**). The difference between theoretical and experimental data can be accounted for by heteroatoms that are expected to be present on the surface of the electrode shifting the voltage and the roughness of quartz capillaries increasing the surface area.

### 2. Mechanism of Adrenaline Oxidation

Electrochemical investigations allow sub-second detection of analytes of interest (29). The method also gives some insight into the electron transport process. To visualize the excitation of molecules under investigation we plotted the redox current against the number of cycles and current in the form of 2-D hotspots (**Figure 2A**). This map allows us to track the electron transfer kinetics of adrenaline. The plot shows an oxidation signature at 0.75 V and a second oxidation signature at 0.9V. The reduction signal at 1.1V (also in Figure 1A) is possibly from the catechol-peroxide interaction. This observation is consistent with our previous recording of dopamine and serotonin on CNT and pyrolytic carbon surfaces (not shown), which shows similar behaviour on oxidative reaction. A reduction signature is seen at -0.2V. From this, it is clear that adrenaline oxidation on CNT-pyrolytic electrodes is not a single step and consists of multiple intermediates.

**Figure 2:**
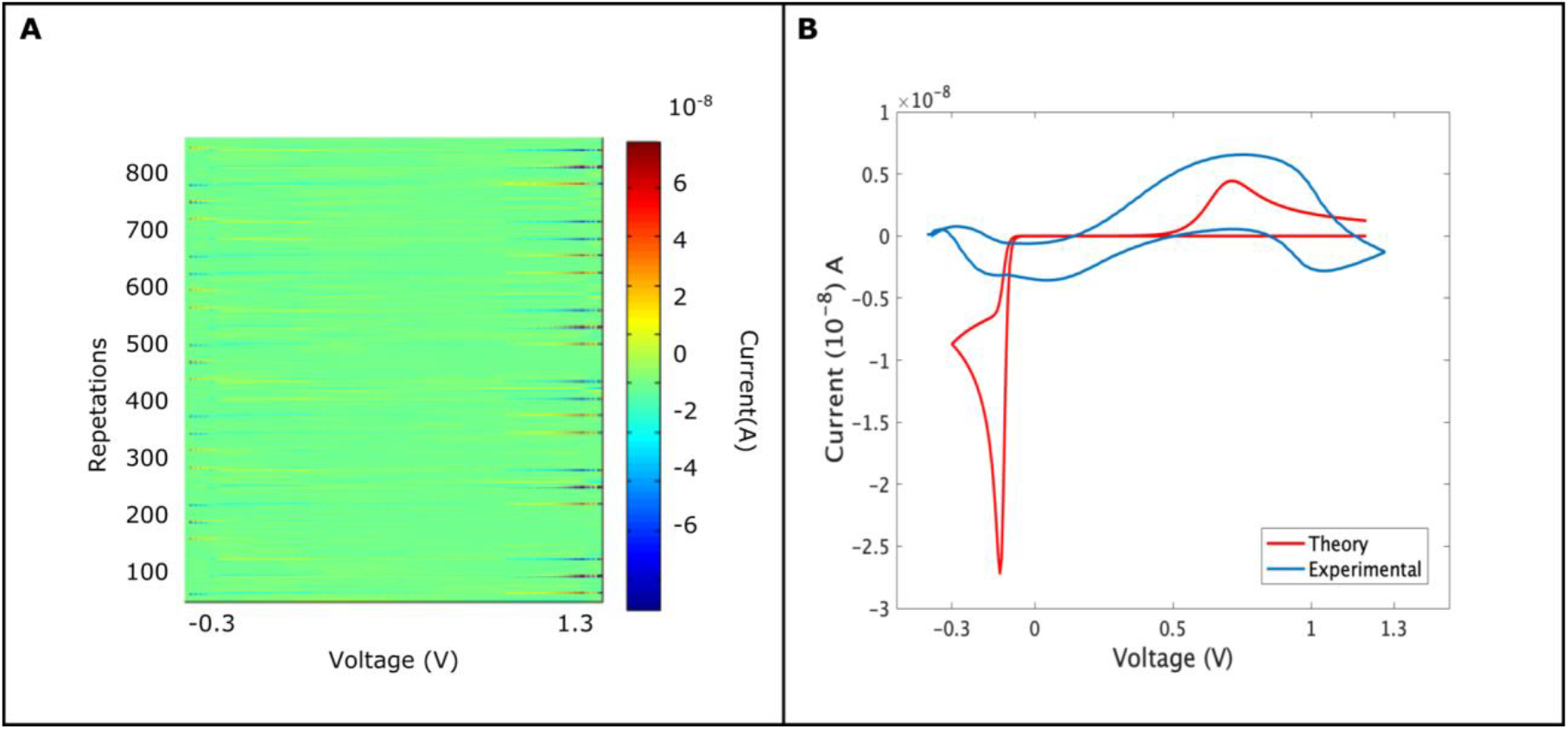
The oxidation and reduction properties of adrenaline using the FSCV protocol. **A**: Shows 2-D hotspot, wherein current is represented on the Y-axis while voltage is represented on X-axis. The time axis is converted into a number of repetitions on the Z-axis. The hotspot shows the oxidation and reduction spots along with the steadiness of current across a number of trials. **B**: Shows the match between experimental and data. Experimental parameters were -0.4V to 1.3V and cycled back to -0.4 V at 600 V s^-1^ repeated at 10 Hz. The simulated data parameters were diffusion constant D 6 × 10^−6^ cm^2^ s^-1^, concentration 1 *µ*M number of electrons transported (n)=2, the rate of electron transport was set to 10^−6^ cm^2^ s^-1^ and chemical rate was 10^−4^ cm^2^ s^-1^.

To explain the electrochemical behaviour of adrenaline on CNT electrodes, we applied the “ECE’’ model presented by Compton’s group (30, 31). A likely mechanism of adrenaline oxidation is the loss of an electron followed by rapid deprotonation. An intermediate radical is formed which loses a further electron to form the quinone product (32).(**Figure 2B)** compares the experimental results with theory. While the peak current for experimental data is higher than the simulation, this is probably because of the rough edges present on the quartz capillaries increasing the effective area. Both peaks are far broader than the simulated ones and the reduction peak shows several maxima, suggesting that a multi stage oxidation and reduction process.

### 3. Selectivity and Specificity

To evaluate if the pyrolytic-CNT electrode is capable of distinguishing analytes, we exposed the electrode to hydrogen peroxide and adenosine. Hydrogen peroxide is known to oxidize at 1.4V on carbon electrodes (5, 6). Background subtracted voltammetry shows the oxidation of hydrogen peroxide on the surface of the electrode (**Figure 3A**) where hydrogen peroxide oxidizes at 0.85 V and has no reduction peak. To understand the electrochemical kinetics of peroxide, we varied the scan rate and measured peak current. The peak current was proportional to scan rate, showing that peroxide is bound to the surface of the electrode (**Figure 3E**). The narrow oxidation peak at 0.85Vshows that oxidation of hydrogen peroxide is rapid, wherein the peroxide is oxidized rapidly into oxygen radical species and water (5, 6). Modelling the reaction shows the oxidation of hydrogen peroxide at 1.3V, using the parameters from (5), however upon increasing the rate of electron transfer we are able to match the theory with experimental (**Figure 3C**).

**Figure 3:**
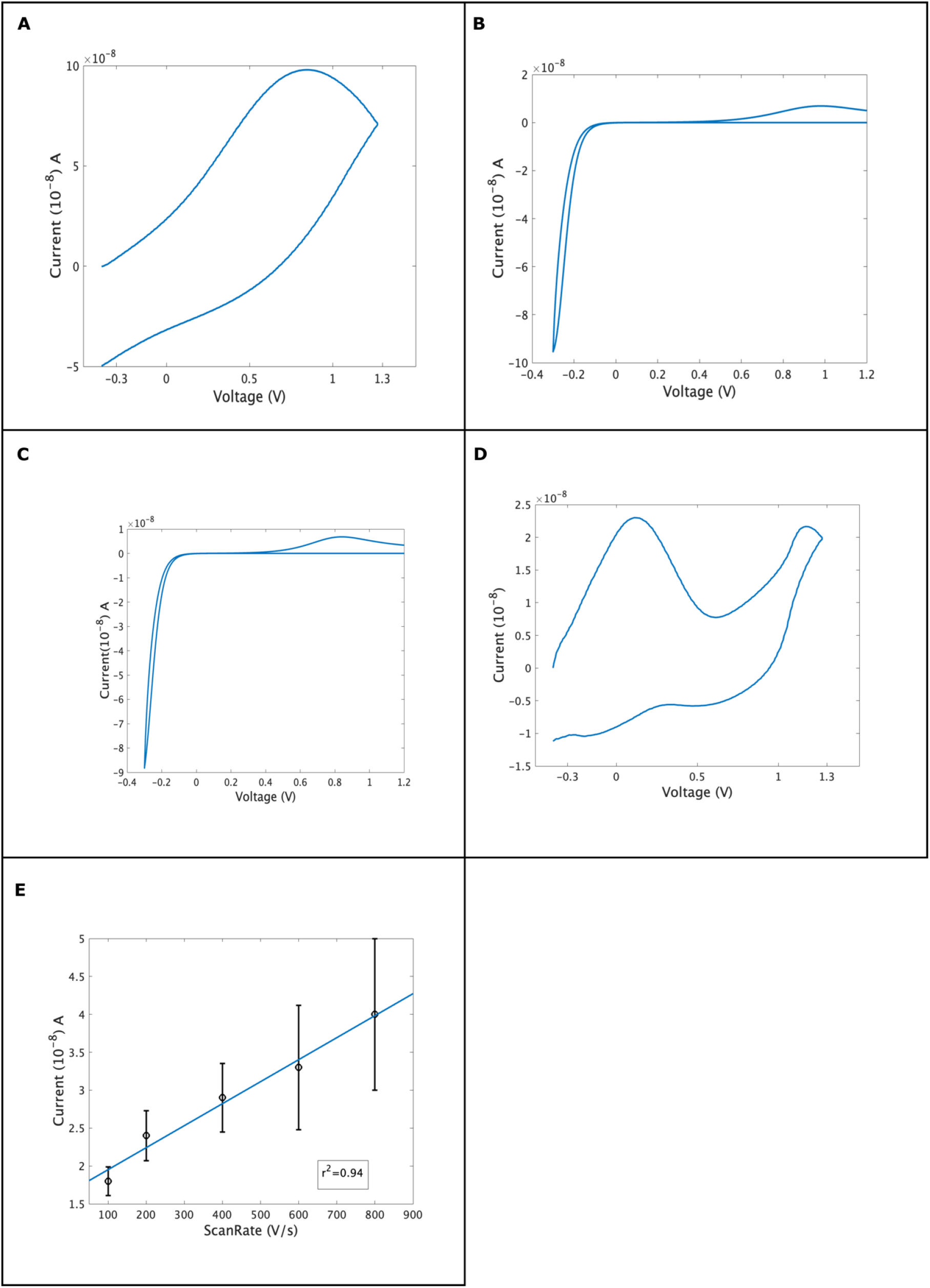
Cyclic voltammogram of 200*μ*M of hydrogen peroxide (H_2_O_2_) on CNT coated electrode. **A:** Hydrogen peroxide oxidizes at 0.85 V. **B**: Electrochemical modelling of reaction. Diffusion coefficient 2.5 × 10^−5^ cm^2^ s^-1^ charge transfer coefficient was 0.5, the number of electron transfer (n)=2 and the electrochemical rate was set to 10^−4^ cm s^-1^ **C**: Increasing the rate of reaction to 10^−6^ cm s^-1^ matches the experimental observation, thus suggesting that CNTs have a faster electron transfer process. **D:** Shows the plot for 1 *μ*M adenosine, which shows an oxidation peak at 0.1 V and a secondary oxidation state at 1.1 V. Broad reduction peaks are seen at 0.7V and 0.2V. **E**: Plot of scan rate *vs* peak current for hydrogen peroxide. The electrode has a sensitivity of 11nA *µ*M^-1^.

Adenosine interferes with the detection of hydrogen peroxide when measurements are conducted using normal carbon electrodes. To investigate the effect of adenosine, we introduced 1*µ*M of adenosine in the flow cell chamber. Adenosine shows a broad oxidation peak at 0.2V and another at 1.2V. Multiple reduction peaks can be observed at 0.6V and 0.2V.(21) (**Figure 3D**). This suggests that pyrolytic carbon electrodes coated with CNTs are able to distinguish between analytes that have a similar oxidation window. Herein, we show that analytes having identical oxidation windows can be distinguished using CNT coatings onto pyrolytic electrodes. Our future work will be setting up a calibration method for multimodal analysis of analytes for detecting the detection limit of our sensor.

## Conclusions

In this work, we showed that carbon nanotibe coated pyrolytic carbon electrodes can be used for the detection of adrenaline, hydrogen peroxide and adenosine and gives different signals for them. The ability of pyrolytic-CNT electrodes to distinguish between analytes by shifting the potential at which they are oxidised coupled with the low overpotential (reducing potential oxidative damage) makes them potentially useful surfaces for the electrochemical detection of neuromodulators. Our future work will include determining the oxidation state of adrenaline on different surfaces using the FSCV method and investigating ways to deconvolute mixed signals as despite the voltage shift of the analytes they are all adsorbed on the electrode and give broad signals so are not fully resolvable yet.

